# Transfer learning for cross-context prediction of protein expression from 5’UTR sequence

**DOI:** 10.1101/2023.03.31.535140

**Authors:** Pierre-Aurélien Gilliot, Thomas E. Gorochowski

**Author notes:** Corresponding author: Thomas E. Gorochowski.

## Abstract

Model-guided DNA sequence design can accelerate the reprogramming of living cells. It allows us to engineer more complex biological systems by removing the need to physically assemble and test each potential design. While mechanistic models of gene expression have seen some success in supporting this goal, data-centric, deep learning-based approaches often provide more accurate predictions. This accuracy, however, comes at a cost — a lack of generalisation across genetic and experimental contexts, which has limited their wider use outside the context in which they were trained. Here, we address this issue by demonstrating how a simple transfer learning procedure can effectively tune a pre-trained deep learning model to predict protein translation rate from 5’ untranslated region sequence (5’UTR) for diverse contexts in *Escherichia coli* using a small number of new measurements. This allows for important model features learnt from expensive massively parallel reporter assays to be easily transferred to new settings. By releasing our trained deep learning model and complementary calibration procedure, this study acts as a starting point for continually refined model-based sequence design that builds on previous knowledge and future experimental efforts.

## INTRODUCTION

When engineers build a system, they typically assume that the parts available to them perform reliable and well-defined functions. This robustness and modularity allows for model-based design of the large and complex systems, as individual parts can be characterised in isolation and the function of their composition with other parts accurately predicted by mathematical models. Synthetic biology has attempted to adopt this approach, with libraries of genetic parts being created and reused across systems ^1^. However, numerous experiment have shown that biological parts often do not behave consistently when used in new ways and often lack the robust modularity that underpins many model-based design approaches ^2^. Multiple factors have been identified as contributing to these contextual effects, including genetic/compositional ^3–5^, functional ^6–8^, host-related ^9–11^, environmental ^12–15^ and experimental ^16^ aspects. Because of this variability in part function, genetic circuit designs often fail to function as expected when first built and typically require lengthy iterative tweaking before a working system is found ^17^.

The 5’ untranslated region (5’UTR) of messenger RNAs (mRNAs) has been found to be particularly prone to contextual effects (especially in prokaryotes), as it contains signals that control translation initiation and can affect overall stability of the mRNA ^18^. This makes the 5’UTR a versatile platform for controlling gene expression by altering sequences upstream of the ribosome binding site (RBS) ^4^, the RBS sequence itself ^19–21^ and possibly synonymously re-coding the N-terminal codons of the protein coding sequence ^22^. The interaction of all these elements results in intricate mRNA interactions and secondary structure formation, which alter protein expression efficiency through various mechanisms including ribosome binding occlusion and pausing ^20;23–25^.

While these contextual effects offer a rich foundation for evolution to shape gene expression, they also make predictive model-guided design difficult because typically genetic parts are only characterised in a limited (often only one) context. Two strategies are therefore typically used to enable effective genetic circuit design. The first consists of mitigating contextual effects by introducing design features that improve the robustness of part function ^26^. For example, adding insulating sequences ^27–29^ and parts (e.g., self cleaving ribozymes) ^1;30;31^ to expression cassettes has proven valuable in reducing compositional effects ^27;31;32^, and at the functional level, feedback control can be implemented to reduce interference between modules ^33–36^. However, the limited number of insulating elements and the additional metabolic cost of implementing feedback compensating mechanisms or tunable systems ^37;38^ means these approaches are not always suitable. A second approach is to not avoid contextual effects, but embrace them by attempting to understand and/or exploit them ^39;40^. One way to do this is by using emerging high-throughput assembly and characterisation approaches ^41–44^ to build large libraries of parts that are then screened to find parts specifically tailored for the task and context at hand. While appealing, difficulties in assembling sufficiently diverse libraries ^45^ and the prohibitive cost of assaying their function, means that these approaches are challenging to use.

In contrast, to better understand contextual effects, mechanistic models have been developed for predicting the effect that some factors have (e.g., the impact of 5’UTR sequence on translation initiation rate). However, these models are often unreliable and again limited to the contextual factors they are able to capture which typically do not extend beyond sequence alone. To overcome these limitations, large genotype-to-phenotype datasets ^41;46;47^ have also been used to employ deep-learning techniques for prediction of part function ^48–50^. However, despite their performance, these models also suffer from poor generalization as they only see data from the biological context used for training. While it would be possible to acquire data across larger numbers of contexts using high-throughput methodologies like Flow-seq ^41;46;47^, and uASPIRE ^48^ to improve model generalization, such experiments are extremely costly, laborious and typically out of reach for most labs ^51^.

A solution to this problem could be to measure a smaller subset of designs that fit within existing experimental workflows (e.g., in a 96-well plate) to provide a smaller set of supplementary training data from the context required. Training a deep learning model from scratch on this small dataset would likely lead to poor performance ^52^. However, a successful technique in computer vision known as transfer learning ^53^ where models are reused across tasks could be used to remove the need to train from scratch. Models like ConvNets ^54^ are typically pre-trained on large datasets such as ImageNet ^55^ and then fine-tuned for specific tasks in which data is scarce. Compared to training using random initialization of the neural network, fine-tuning of a pre-trained model offer superior performance while requiring significantly fewer data. Transfer learning applied to deep learning models of sequence-to-function relationships could therefore have the potential to create many tailored models for each context to be used during design.

In this work, we explore the possibility of using transfer learning to adjust the weights of a pre-trained deep learning model able to predict protein expression from 5’UTR sequence in *E. coli* such that it enables us to make accurate predictions in new contexts (**Figure 1**). We demonstrate the effectiveness of a hybrid neural network architecture to model the effects of base pair mutations in the RBS sequence and show that the features learned during training can be adapted to different sequence and experimental contexts using a simple transfer-learning procedure. Using this approach, we then quantify the amount of data needed to successfully transfer models across a wide range of different contexts to identify the appropriate data calibration procedure. Our pre-trained model and simple fine-tuning pipeline are a valuable tool to support model-based 5’UTR engineering, and demonstrate that transfer learning may provide a means for applying data-centric models in biological design more broadly.

**Figure 1:**
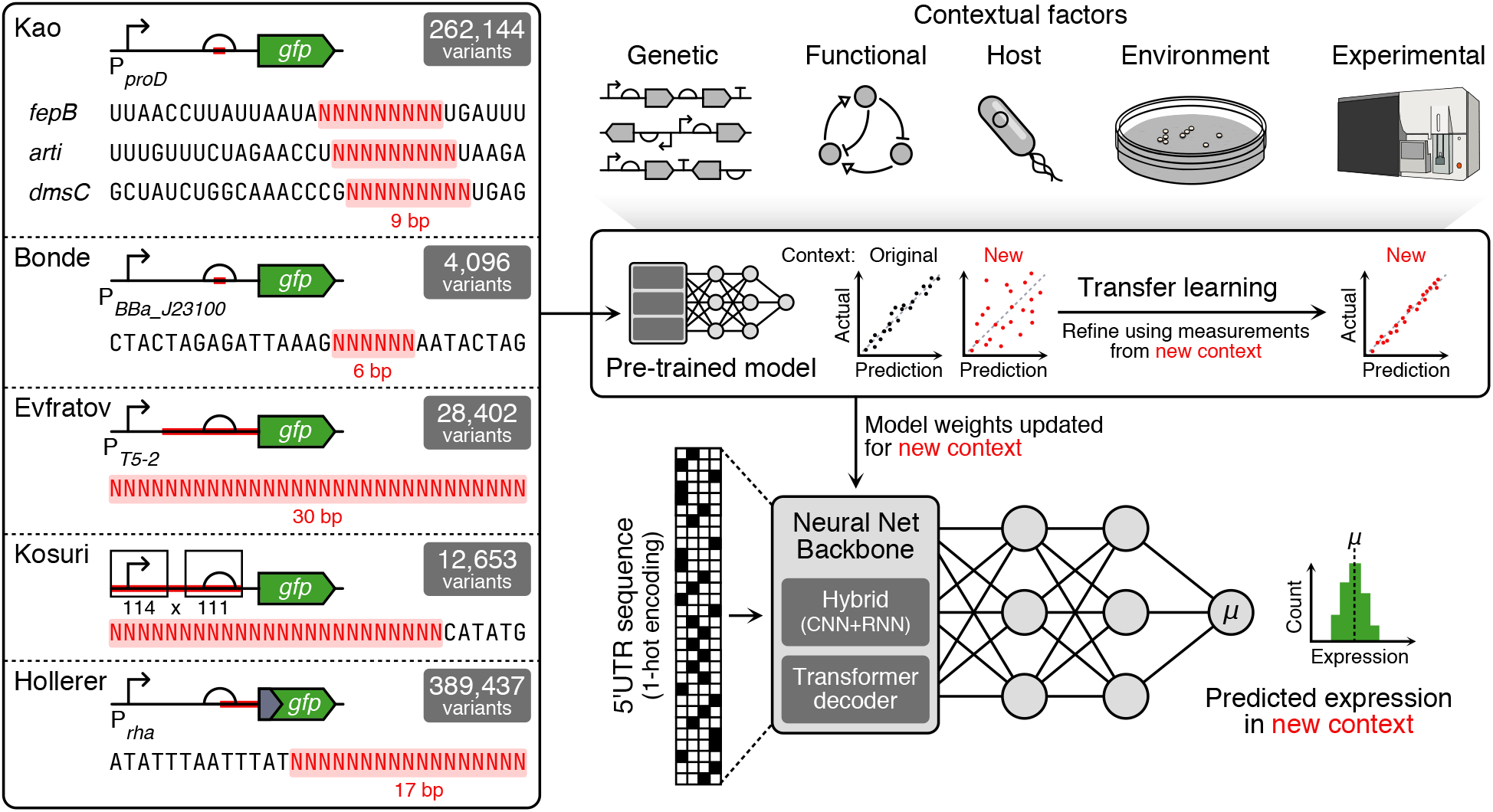
Transfer learning pipeline to adapt pre-trained models for new contexts. Left box contains the design of the datasets considered to pre-train and test the models developed. Randomised sections of the 5’UTR are shown in red. On the right-hand side, top elements indicate the various contextual factors that can affect the function of a genetic part. Middle box shows the workflow we use, starting with a pre-trained deep neural network and then fine-tuning this model via a transfer learning approach with a small number of samples from a new context in which predictions need to be made. Lower region shows the general structure of the different neural networks considered (either Conv-LSTM or the frozen transformer decoder backbone from the RNA foundational model), which predict mean protein expression from a 5’UTR sequence.

## RESULTS

### Predicting translation initiation rate from 5’UTR sequence using deep learning

We began by developing a deep learning based model to predict protein expression rate from the 5’UTR sequence. Deep learning models are ideally suited for modelling sequence-to-function relationships due to their ability to learn complex nonlinear mappings to any level of precision. Given their high-levels of performance in genomic studies ^56^, we investigated the ability of Convolutional neural networks (CNN) and hybrid Convolutional-Long Short Term Memory (Conv-LSTM) neural networks to accurately learn the log-fluorescence mean from 5’UTR sequence alone. Both models were initially trained on a dataset generated by Kuo and colleagues ^19^, where fixed 5’UTR sequences contains a 9-nt variable Shine Dalgarno (SD) region (**Figure 1**). We specifically used the *fepB* subset when developing our models. To ensure high-quality data was used for training, the FORECAST package ^51^ was used to fit normal distributions to the log-fluorescence data using maximum likelihood estimation. This inference step returned for each 5’UTR sequence the mean (*µ*) and standard deviation (*σ*) of the underlying log-normal fluorescence distribution, which has been shown to be a better representation when comparing gene expression than using raw fluorescence data ^57^.

Rather than defining a fixed architecture for our models, we used neural network hyperparameter optimization to find suitable designs that maximised the accuracy of predictions (**Methods**). This revealed that both types of model could reach a similar level of performance, with a Mean Squared Error (MSE) = 0.48 for the *fepB* dataset. Interestingly, the Conv-LSTM model achieved this performance with eight times fewer parameters than the CNN model, having 1.1 million versus 8.7 million parameters, respectively (**Figure 2a**). The efficiency of the Conv-LSTM model is likely due to how the convolutional and recurrent layers capture complementary aspects of the biophysical gene expression process. CNNs excel at extracting motifs/patterns in sequences, which here are known to play an important role (e.g., facilitating base pairing between the mRNA and the 3’end of the 16S ribosomal RNA). In contrast, the LSTM module is able to efficiently model the ordering and distance between such motifs, which again is known to be critical for controlling translation initiation rate ^20^. This allows the Conv-LSTM model to exploit patterns in both of these features more efficiently than the CNN can alone. The best performing Conv-LSTM architecture was found to achieving a median absolute error = 0.36 on the test set (**Figure 2b**). This is equal to the minimum achievable error for this dataset, computed by considering the median width of the 99.7% confidence interval for the log-fluorescence mean (**Supplementary Figure 1**). Practically, this corresponds to a median absolute percentage error of 35% in the raw fluorescence data (**Figure 2b**). Curiously, the perceptron block of the Conv-LSTM model selected after hyperparameter optimization contained only a single intermediate layer. This suggests a relatively simple relationship between the features extracted by the backbone (CNN and LSTM units) and the output fluorescence (**Figure 3a**).

**Figure 2:**
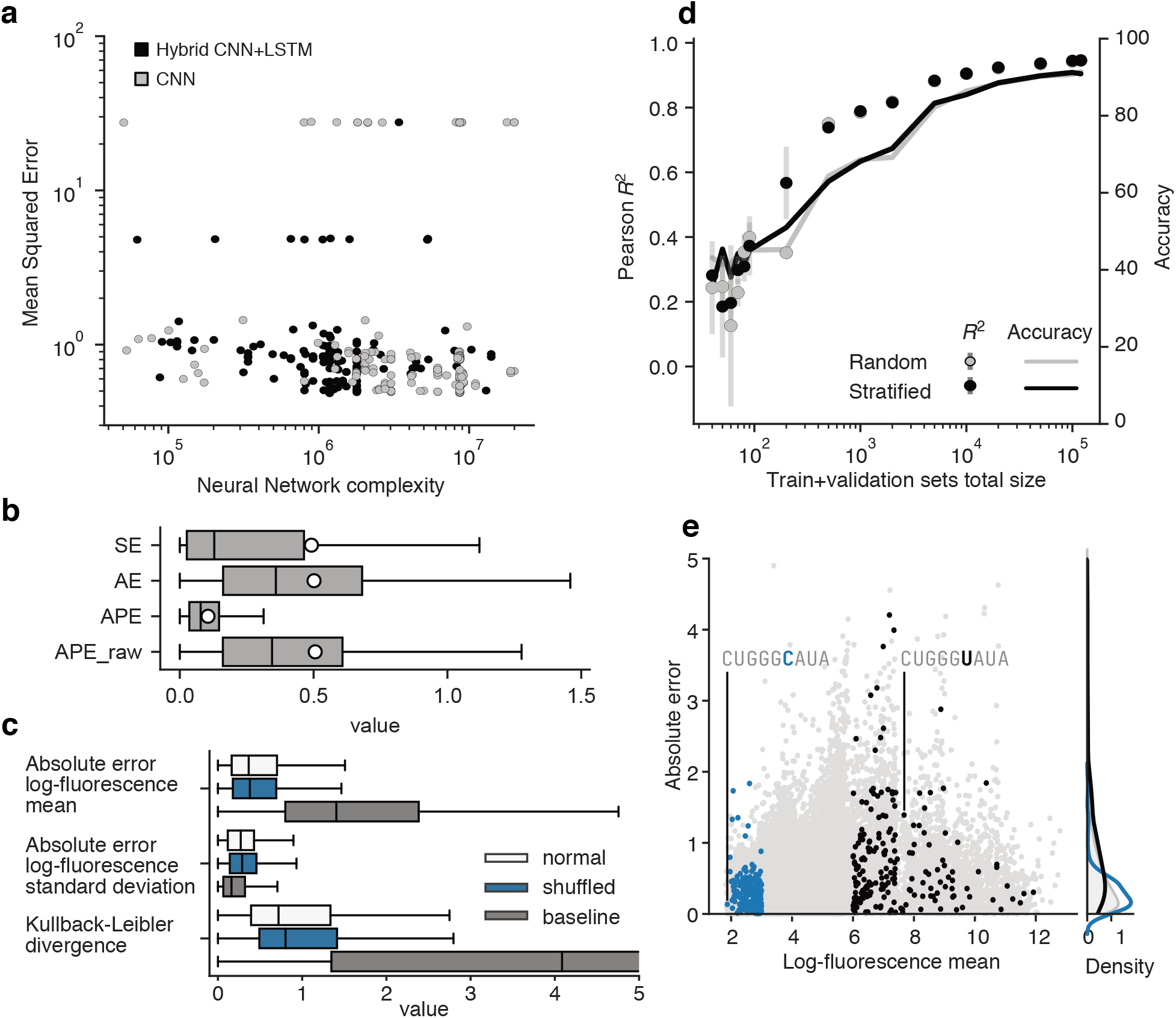
Deep learning accurately predict protein expression from 5’UTR sequence. (**a**) Number of parameters and best mean squared error (evaluated on the validation set) of all models trained during the hyperparameter optimisation phase using the *fepB* dataset. (**b**) Distribution of the test set performance metrics (SE: Squared Error, AE: Absolute Error, APE: Absolute Percentage Error of the log-fluorescence data, APE raw: Absolute Percentage Error of the fluorescence data) for the Conv-LSTM model trained to predict the mean log-fluorescence from 5’UTR sequence. (**c**) Evaluation of the performance of models trained to simultaneously predict the log-fluorescence mean and standard deviation. (**d**) Performance of the best Conv-LSTM model predicting the log-fluorescence mean using varying size and composition of training data.Error bars indicate the standard deviation observed when training three models using different random seeds. (**e**) Distribution of prediction errors for the test dataset. Activity cliffs (pairs of sequences with highly different 5’UTR strength that differing by only a single base mutation) are colored in blue and black. Each sequence in blue has a corresponding sequence in black. Two representative sequences are labelled with the mutation highlighted.

**Figure 3:**
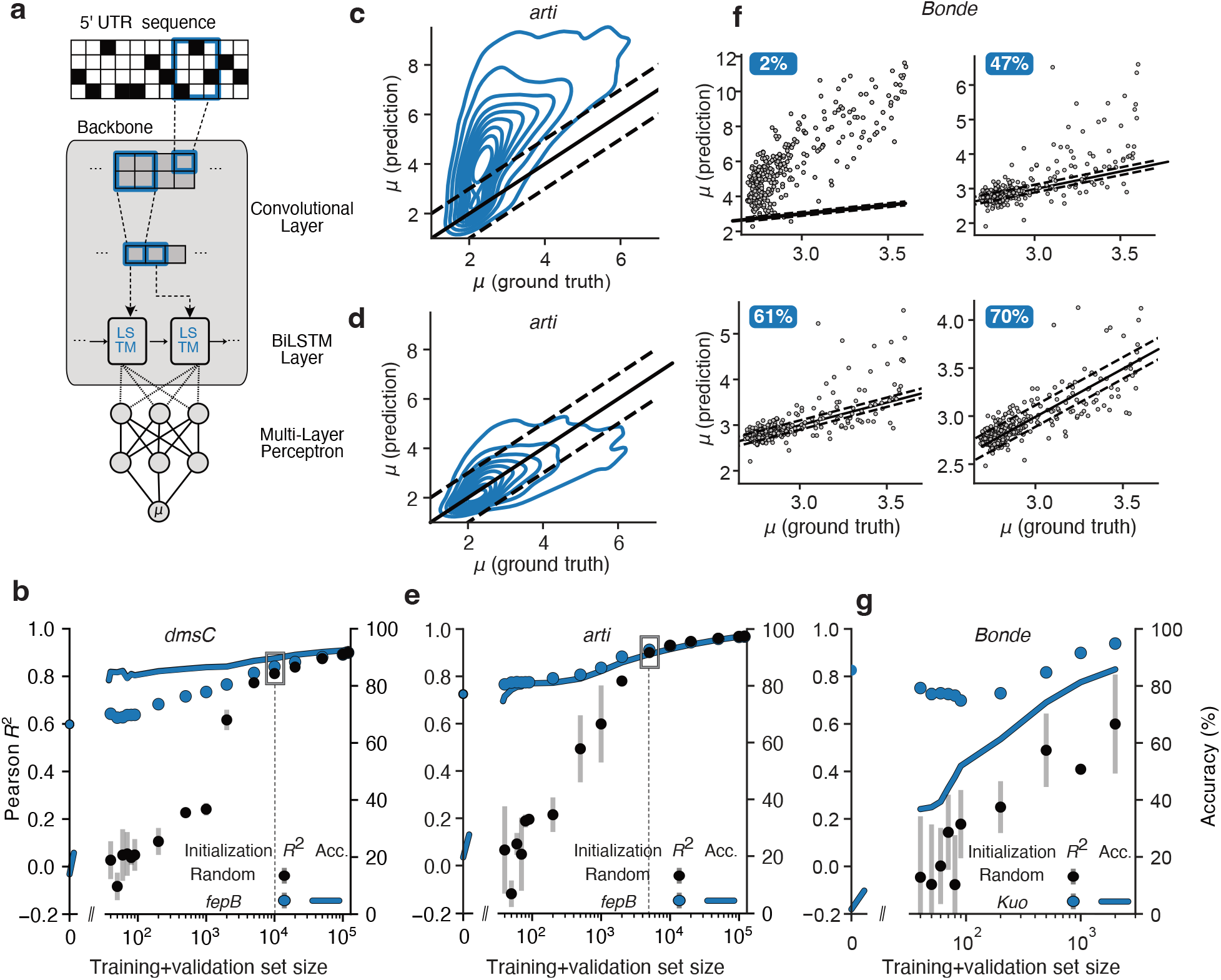
Adaptation to new genetic and experimental contexts using transfer learning. (**a**) Architecture of the best performing neural network predicting 5’UTR strength. Weights across all layers were adjusted during fine-tuning to different genetic and experimental contexts. (**b**) Test set (*dmsC* data) performance of the fine-tuned Conv-LSTM model (pre-trained on *fepB*)) versus a Conv-LSTM model trained from scratch (Random initialisation). The right *y*-axis represents accuracy and the box shows when both models are of similar performance. Error bars indicate the standard deviation observed when training three models using different random seeds. (**c**) Kernel density estimates of the test set (*arti* data) predictions (mean log-fluorescence) of the Conv-LSTM model trained on *fepB*), before fine-tuning. (**d**) Kernel density estimates of the test set (*arti*) predictions (mean log-fluorescence) of the Conv-LSTM model trained on *fepB*), after fine-tuning using 90 sequences. Predictions were averaged across three models trained using different random seeds. For (**c**) and (**d**), the black line shows *x* = *y* (i.e., perfect estimation) and the dashed black lines indicates the *±*1.7 margin around it. Iso-lines are plotted in blue, with the outermost containing 90% of the data. (**e**) Test set (*arti* data) performance of the fine-tuned Conv-LSTM model (pre-trained on *fepB*)) versus a Conv-LSTM model trained from scratch (Random initialisation). Left *y*-axis indicates the Pearson *R*^2^ value, right *y*-axis indicates the accuracy and the box shows when both models are of similar performance. Error bars indicate the standard deviation observed when training three models using different random seeds (**f**) Test set (Bonde data) predictions (mean log-fluorescence) of the fine-tuned Conv-LSTM model (trained on all Kuo data) fine-tuned using 0 (top left), 90 (top right), 200 (bottom left), 500 (bottom right) sequences. Predictions are on the *y*-axis, and ground truth is on the *x*-axis. Black line shows *y* = *x* (i.e., perfect estimation) and dashed black lines indicates the *±*0.2 margin around it. Accuracy is indicated on the top left for each graph. (**g**) Test set (Bonde data) performance of the fine-tuned Conv-LSTM model (pre-trained on all Kuo data) versus a Conv-LSTM model trained from scratch (Random initialisation). Left *y*-axis indicates the Pearson *R*^2^ value. Error bars indicate the standard deviation observed when training three models using different random seeds.

The Flow-seq experiments used to generate the Kuo dataset involves the sorting of genetic variants into several contiguous bins, therefore capturing varying levels of output fluorescence. This allows for higherorder statistical information to be measured such as the fluorescence standard deviation by analysing the spread of genetic variants across the bins. Such information is indicative of the level of variability or noise within the cell and across the population and potential useful for developing parts to control the variation in gene expression. We explored whether it was possible for our deep learning model to learn the standard deviation of the fluorescence output for each 5’UTR. To assess whether our model could be used to predict this feature, the original Conv-LSTM model was modified to include an additional neuron in the output layer to predict the standard deviation of the log-fluorescence. This new head enables the model to learn a distribution by minimizing the distance with the ground truth-distribution, measured using the Kullback Leibler (KL) divergence ^58^. After optimizing the hyperparameters of this new architecture, we assessed the predictions of two instances of this model, either trained using the original dataset or using an altered version of this dataset where the standard deviation information was shuffled between samples.

We found that the difference in Mean Absolute Error (MAE) when predicting the mean log-fluorescence *µ* for both models was not statistically detectable (*<*0.005, p-value *>* 0.1, Welsh t-test) **(Figure 2c**). When predicting the standard deviation *σ* of the log-fluorescence, the model trained on the shuffled dataset performed slightly worse, with a MAE = 0.33 versus 0.31 for the model trained on the normal dataset. However, this difference is not practically significant, as the median 99.7% confidence intervals width for *σ* is 10 times bigger (approximately 0.31; **Supplementary Figure 1**). The similarity between errors suggests that either the model is not well-suited to learning the standard deviation, or the experimental data does not contain sufficient information for estimating this parameter. Indeed, had the data contained a learnable signal, the performance of the model trained on the original data would have been higher than the model trained on shuffled data. Additionally, the model trained on the original data was found to perform worse than a simple baseline model which always returns the average log-fluorescence standard deviation. Such baseline model leads to a MAE = 0.16 vs 0.33 for the model trained on the normal dataset **Figure 2c**). It is unlikely that the issue stems from the inability of deep learning model to model the data, as we allowed for the neural network to expand its capacity during the hyperparameter optimization. It is therefore more likely that unavoidable noise and biases in the experiments used to collect the data is the cause ^48^. These results suggested that the Conv-LSTM model is not able to find useful patterns in the data for accurately predicting the standard deviation of the fluorescence measurements, and so we focused our efforts on prediction of the mean log-fluorescence.

As high-throughput assays are expensive and time-consuming, we wanted to better understand the minimal number of samples required to train models across a range of contexts. To do this, we performed a data-efficiency analysis of the Conv-LSTM model where we trained models from scratch on sub-sets of the *fepB* training and validation data. We found a consistent increase in performance as the number of training examples grew, with the Pearson’s *R*^2^ increasing from 0.2 to 0.94 (**Figure 2d**). Because the Pearson’ *R*^2^ depends on the value and distribution of the response variable (here the test set values), we also used an accuracy metric to account for the non-standardized units used to measure fluorescence during each experiment. The metric considers the percentage of predictions within a specified margin of the ground truth. Here, we chose the value of this margin to be an eighth of the data spread (for the *fepB* dataset the margin = 1.12). Based on this metric, we found that accuracy also increased with the number of examples, and like the Pearson’s *R*^2^, exhibited diminishing returns. This highlights a trade-off between prediction quality and the size of the training dataset. By assuming a cost function that logarithmically grows with the number of constructs characterized (typical for high-throughput experiments), the elbow point optimizing this trade-off was located at 5,000 samples, corresponding to a Pearson’s *R*^2^ = 0.74 and an accuracy of 82% (**Figure 2d**). We also assessed whether the composition of the training set sequences had an influence on the performance reached and compared the performance when selecting sequences by using stratified sampling with respect to the distribution of 5’UTR strengths. We found that performance was similar regardless of the type of sampling used, with differences between *R*^2^ not practically nor statistically significant (**Supplementary Figure 2**).

Modifying the 5’UTR is a popular approach for controlling gene expression in prokaryotic cells ^59;60^ because small changes to the sequence can dramatically alter protein expression level ^19;21;61^. Not only is the Conv-LSTM model able to accurately predict those changes on average across the test sequences, as shown by the MAE = 0.5 (**Figure 1b**), but it is also able to distinguish between strong and weak 5’UTRs that differ by only one nucleotide. Indeed, the model had a MAE = 0.3 and 0.8 for the weak and strong expression test sets, respectively, with both of these being smaller than the accuracy margin of 1.12. With the model performing well on the test set, the ability to resolve activity cliffs (pairs of 5’UTR with highly different protein expression strengths that differing by only a single base mutation), provides evidence that our Conv-LSTM model does not over-fit to the training set, but is instead able to generalize across the entire 5’UTR landscape. Such nucleotide level sensitivity is difficult to obtain with non-deep learning models, and we typically observed highly similar predictions for the activity-cliff sequences when using the biophysical models. Specifically for the RBS calculator (v2.1) ^21^, we found that the weak-strong ordering for each pair was incorrect 30% of the time, with the ratio between the maximum translation initiation rates never exceeding 5 for 95% of the pairs **Supplementary Figure 6**). These results show that we can accurately predict how the mutations within the RBS region of a 5’UTR sequence affect protein expression using a neural network combining both convolutional and recurrent layers. Although the model is close to optimal, this guarantee only holds for the specific distribution of the *fepB* dataset. It is therefore essential to evaluate the performance of this model and its transferability across different genetic contexts, host strains and experimental condition to see how well it truly captures all the factors essential for accurate prediction.

### Transfer learning to different sequence contexts

High-throughput genotype-to-phenotype experiments based on methods like Flow-seq, often screen libraries of sequences harbouring a constrained design (e.g., a large sequence containing a small variable region). In the Kuo dataset, the SD sequence within the RBS is a 9-nt variable region, flanked by different fixed sequences corresponding to different genetic contexts (i.e., *fepB, arti* and *dmsC*). Differences in the nonvariable regions can have a strong influence on protein expression through different local mRNA interactions and folding ^23;62–64^ and become confounding effects during the training of a neural network. Although our first Conv-LSTM model was exclusively trained on the *fepB* context of the Kuo dataset, the features learnt might still be meaningful in different 5’UTR contexts, but some tuning of the network may be required.

Transfer learning allows features learnt by a model to be adapted for use in a different context by adjusting the neural network parameters after processing a limited number of samples from the new context ^53^. To see whether such an approach might work in our case, we investigated how efficient a simple fine-tuning procedure was by starting with our pre-trained Conv-LSTM for the *fepB* context, and then testing on other contexts after adjusting the weights using a small number of new training examples. To quantify the data requirements of this transfer leaning approach, we considered the *dmsC* and *arti* datasets from the Kuo experiment (**Figure 1**). For each of these, the data was collected in the same way as for the *fepB* dataset. The only differnce being the fixed regions of the 5’UTR sequence. To assess the improvement that transfer learning provided, the Conv-LSTM model with pre-trained weights was trained alongside a model with the same architecture but random initial parameter (i.e., weights). We found that while both models converged to a similar accuracy after 10,000 training examples were provided, the pre-trained model had significantly better initial performance, with Pearson’s *R*^2^ = 0.73 for the *arti* dataset and *R*^2^ = 0.6 for the *dmsC* dataset before fine-tuning (**Figure 3b,e**). This initial performance was also superior to several recent RBS strength predictors such as RBSeval (*R*^2^ = 0.58 for *arti* and *R*^2^ = 0.47 for *dmsC*) and EMOPEC (*R*^2^ = 0.5 for *arti* and *R*^2^ = 0.47 for *dmsC*) (**Supplementary Tables 2 and 3**). This suggests that these models are not effective at predicting small SD mutations. A possible explanation for this behaviour is that the models rely on RNA secondary structure predictions, which might not adequately capture the effects of small mutations within the SD sequence. This hypothesis is supported by the poor correlation between protein levels and SD:aSD base pairing energy (*R*^2^ = -0.5 for *arti* and *R*^2^ = -0.42 for *dmsC*)and mRNA folding energy (*R*^2^ = 4 *×*10^*−*3^ for *arti* and *R*^2^ = 0.01 for *dmsC*) (**Supplementary Tables 2 and 3**).

As expected, the performance of fine-tuned models steadily improved with the number of new training examples provided (**Figure 3b,3e**). However, the improvements in Pearson’s *R*^2^ become small after 90 examples for fine-tuning (≈0.05 for both the *dmsC* and *arti* datasets) suggesting that the model has already been refined sufficiently to the new context with the accuracy jumping from 14% to 84% for the *dmsC* dataset, and from 23% to 85% for the *arti* dataset (**Methods**). Fine-tuning also regularizes training when data is limited (*<*2,000 samples), as many neural networks trained from scratch often predicts a constant value, the mean log-fluorescence, instead of learning meaningful patterns (**Supplementary Figure 4**). This training instability is evident by the large standard deviation observed when training different models using different seeds (**Figure 3b,e**). Having such behaviour is undesirable as it hinders the use of commonly used techniques for improving model performance, like bagging, which involves averaging predictions from a group of independent models ^65^.

In addition to randomly selecting 5’UTR sequences from a new context for fine-tuning, we also assessed if selective sampling might accelerate the transfer learning process. We found that stratified sampling of the data from the new context based on fluorescence level did not significantly increase the performance of the fine-tuned model (*p >*0.2, Welch’s t-tests for equal Pearson correlation between stratified vs random sequence sampling (**Supplementary Figure 3**).

### Transfer learning to new sequence and experimental contexts

Transcription and translation are physically coupled in prokaryotes ^66;67^ allowing for regions of the promoter sequence downstream of the transcription start site to impact translational processes ^4;62^. As a result, the promoter sequence could become a confounding factor in high-throughput experiments where typically only a single promoter is used to drive transcription of a reporter gene.

To explore whether our fine-tuning approach could adapt to changes in promoter context, we made use of another dataset collected by Bonde and colleagues ^68^, where they characterized all mutations of a 6 nucleotide-long SD sequence in the RBS of the 5’UTR. Compared to the Kuo dataset on which our model was pre-trained, the Bonde dataset uses a different promoter (P_*proD*_ versus P_*BBa J23100*_, respectively), has a different fixed 5’UTR context around the RBS, and the reporter protein (GFP) sequence has a different codon usage bias. Furthermore, differences in the experimental setup (i.e., the FACS machine used to measure fluorescence) meant that the fluorescence units used to report the protein expression strength also differed from the Kuo dataset with the Bonde test set taking values in the interval [2.7, 3.6].

We began by retraining the Conv-LSTM model using all data from the Kuo dataset (using all three 5’UTR contexts). This increased the models’ exposure to different genetic contexts around the SD sequence. Fine-tuning this new model on the Bonde dataset revealed a large improvement in performance, with the accuracy jumping from 2% to 47% after fine-tuning with a set of only 90 examples (**Figure 3f,g**). In contrast, the performance of the model trained from scratch is always inferior (**Figure 3g**). The Bonde dataset lacks sufficient data to determine when the performance of these two models becomes comparable, which is consistent with the previous analysis of the Kuo dataset, which identified such a point between 10,000 and 20,000 sequences (**Figure 3b,e**). We found that incorrect predictions from the fine-tuned model initially tended to be large positive deviations from the ground truth. These differences were corrected as the number of fine-tuning examples increased (**Figure 3f**). These correspond to strong RBSs, which are less frequent and therefore require more fine-tuning data to observe them. The steady increase in accuracy for the fine-tuned model highlighted the ability of the model to both adapt to a different genetic context and measurement units. This need to concurrently adapt to two differing demands may explain the initial drop in *R*^2^ when the number of fine-tuning examples is low (**Figure 3g**).

To understand the importance of the backbone module (i.e., the Convolutional and LSTM layers) of the neural network, we conducted a variant of fine-tuning in which the backbone parameters were frozen and only the parameters of the MLP layers could be updated. We hypothesised that the backbone may capture sufficient features that only modifications in the way these features are used by the final MLP layer, might be sufficient to speed up the transfer learning process. We found this to not be the case, with the alternative fine-tuning procedure requiring at least 2,000 sequences to achieve an accuracy of 40% (**Supplementary Figure 6**), highlighting the importance of modifying parameters of the entire neural network. This suggests that, in addition to the conversion of fluorescent measurement units between data sets (i.e., scaling), which modifications to the final MLP layer could accommodate, new features are potentially learned during the fine-tuning process.

### Transfer learning to diverse global 5’UTR contexts

So far we have trained the Conv-LSTM model on data sets that include a small variable sequence within the 5’UTR. The lack of diversity across the full length of the 5’UTR could lead to issues in applying the model in broader contexts. To address this, we examined whether the Conv-LSTM model was capable of adapting to 5’UTR sequences that differ across their full length. We made use of a dataset collected by Kosuri and colleagues ^4^, which measured gene expression for a fully combinatorial library of 114 promoters and 111 ribosome binding sites. After extracting the mean log-fluorescence from the raw sequencing data and filtering out low quality sequences (**Methods**), we were left with a collection of 63 promoters with varying numbers of associated RBSs. Each promoter sub-dataset was then divided into three sets: a training set (for fine-tuning, whose size varied from 12 to 61 sequences), a validation set (for fine-tuning early stopping, always consisting of 20 examples) and a test set (for evaluation, always consisting of 30 examples to minimize sampling variations).

We began by fine-tuning the Conv-LSTM model pre-trained on the Kuo dataset using data for each separate promoter from the Kosuri dataset. Despite the small amount of training data, the benefits from fine-tuning were still apparent, as demonstrated by the 40% increase in accuracy obtained for the apFAB80 promoter whose training set consisted of only 16 sequences (**Figure 4a,b**). Overall, for all promoters, finetuning improved the accuracy of the predictions by 13.5% (*p <* 3.3 *×* 10^*−*6^, paired permutation testing of equal mean after and before fine-tuning) (**Figure 4c**). However, the improvements were not uniform across all promoters with some displaying poor accuracy even after fine-tuning. Ignoring the issue of limited numbers of samples, this suggests that the features learnt during training on the full Kuo dataset were not insufficient to capture all of the biological phenomena observed for every promoter in the Kosuri dataset. This is supported by the negative transfer observed for 12.3% of the promoters (from promoter 0 to promoter 7; **Figure 4c**), although the differences were not statistically significant (all *p*-values *<*0.75 using paired permutation tests for equal mean after and before fine-tuning for each promoter) (**Supplementary Figure 7**).

**Figure 4:**
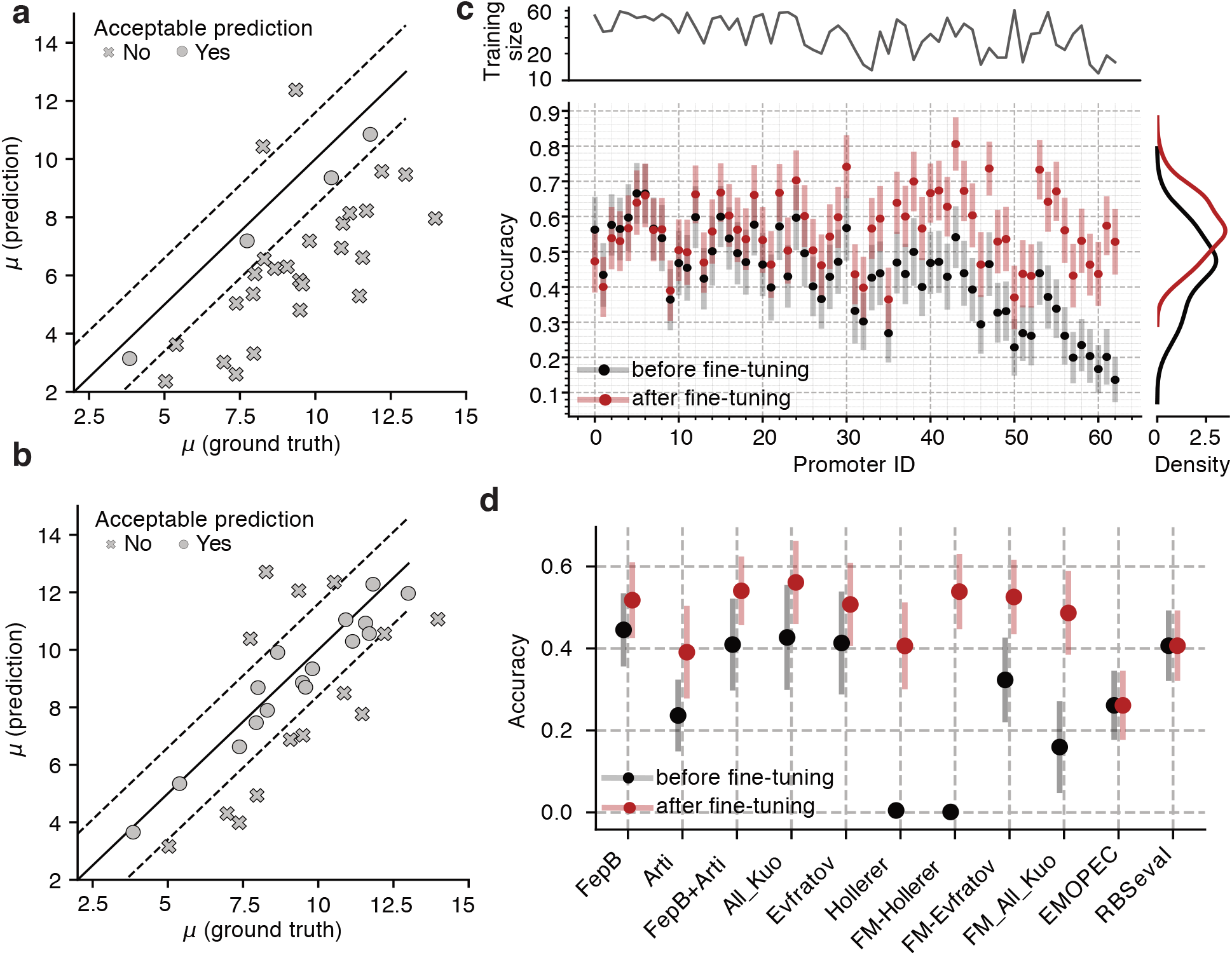
Transfer learning accomodates for gloablly different sequences with only a handful of sequences. (**a**) Test set predictions of the mean log-fluorescence (Kosuri dataset, promoter apFAB80) using the Conv-LSTM model pre-trained on all Kuo data, before fine-tuning. (**b**) Test set predictions of the mean log-fluorescence (Kosuri dataset, promoter apFAB80) using the Conv-LSTM model pre-trained on all Kuo data, after fine-tuning using only 16 examples in the training set and 20 examples in the validation set. (**c**) Accuracy of the Conv-LSTM model pre-trained on all Kuo data, tested on each Kosuri promoter sub-dataset, before and after fine-tuning. Number of sequences used in the training set for each promoter sub-dataset are reported. To increase visibility, promoters were ranked according to their accuracy ratio after and before. Standard errors were estimated using 500 bootstrap samples. (**d**) Mean accuracy across all Kosuri data before and after fine-tuning for every model considered. A prefix FM indicates that the embedding from the RNA foundational model is used, otherwise, one-hot encoding of the 5’UTR is used.

To assess the amount of information learnt by the Conv-LSTM model, we compared its performance to a collection of additional models pre-trained on different datasets. First, to test the importance of highly diverse 5’UTR data for learning generalizable features, we compared the performance of the Conv-LSTM model trained on different combinations of 5’UTR contexts from the Kuo dataset. Models trained on a unique 5’UTR contexts (‘FepB’ or ‘Arti’) had lower accuracies than models trained on combined 5’UTR contexts (‘FepB+Arti’ or ‘All Kuo’), with the model trained on all the 5’UTR contexts (‘All Kuo’) achieving the best mean accuracy of 56% across all promoter datasets (**Figure 4c**). For comparison, this accuracy was found to be higher than several widely used RBS strength predictors, with EMOPEC ^68^ and RBSeval ^69^ achieving accuracies of 26% and 40%, respectively (**Figure 4c**), which further highlights their limited performance. Interestingly, we found that RBSeval sometimes failed to provide useful predictions at all, returning almost identical values for some promoter subsets (**Supplementary Figure 8**).

Increasing the diversity of the 5’UTR dataset can also be achieved by training the deep learning model on globally diverse sequences, which are likely to cover a wider range of biological factors/processes that could influence gene expression. To test this, we trained two new Conv-LSTM models on two new datasets where the 5’UTR composition was globally varied (the Evfratov ^64^ and Hollerer ^48^ datasets; **Figure 1**). This increase in sequence diversity did not lead to any significant improvement in model performance, with a lower mean accuracy for the Evfratov-trained model (50%, *p* = 1.2 *×* 10^*−*3^, paired permutation test for equal mean) and for the Hollerer-trained model (41%, *p* = 3.3 *×* 10^*−*6^, paired permutation test for equal mean). Remarkably, the limited fine-tuning examples were sufficient to adapt between the different experimental setups (i.e., fluorescence measurement units) used in the Hollerer dataset which covered the interval [0, 1] (measuring the area under the flipping ratio curve of a recombinase reporter system ^48^) and the Kosuri dataset which covered the range [2, 15] a.u. (measured using a fluorescence-based assay). These results suggest that that the biological phenomena observed when mutating the SD region and its local vicinity is sufficient for sequence models to extract useful generalizable patterns related to protein expression.

A further question we posed was whether fine-tuning could be improved by using additional biological information. To that end, we utilized embeddings computed by an RNA foundational model (FM) ^70^ instead of the raw 5’UTR one-hot encoding. The RNA-FM was trained through self-supervised learning on 23 million non-coding RNA sequences, which consisted of reconstructing for each RNA sequence the identity of a few (15%) masked nucleotides. The underlying transformer model makes use of the selfattention mechanism ^71^ to ensure the model learns contextual dependencies. Earlier work demonstrated the higher information content of these embeddings as input for a convolutional neural networks on a large class of structure and function prediction tasks, including eukaryotic 5’UTR translational strength prediction ^70^. Here we used their architecture and retrained their model on all the libraries considered so far, before fine-tuning it on the Kosuri dataset.

Models trained using this embedding performed differently depending on the type of variation in the 5’UTR sequences. When trained on the Kuo dataset, where variation in sequences is limited to a 9 nt region, using these embeddings as input rather than the one-hot encoding resulted in less ability to generalize, with a decrease of 27% in mean accuracy for the Kosuri dataset. Fine-tuning was also found to be less efficient, with a significant decrease in mean accuracy of 7% (*p* = 1.6 *×* 10^*−*5^, paired permutation test for equal mean) (**Figure 4d**). In contrast, using the RNA-FM embeddings for models pre-trained on globally different 5’UTR sequences (i.e., Evfratov and Hollerer datasets) led to similar or better fine-tuning performance, with a significant increase in mean accuracy of 13% after fine tuning for the Hollerer-trained model (*p* of *×* 10^*−*6^ paired permutation test). This suggests such embeddings may be appropriate for sequences with a high degree of global diversity. Because the sequences from the Kuo dataset only vary by one-nucleotide, the embeddings computed by the RNA foundational model might attenuate this fine-grain information.

## DISCUSSION

The ability to precisely control gene expression is crucial for the development of complex genetic circuitry and the reprogramming of cells. Designing 5’UTRs is a popular approach to achieve this goal, but often requires the screening of many variants in order to find a sequence with precisely the desired function. In this work, we have shown that deep learning models of 5’UTR strength can be derived from high-throughput experimental data and that they outperform many classical methods to predicting protein expression rate from 5’UTR sequence. Our analysis suggests that current high-throughput assays provide sufficient information to accurately predict mean protein expression, but that higher order statistical information like the variance are not well captured. Furthermore, we have shown that the design of a Flow-seq experiment can biases the representations learned by a neural network, limiting the ability to use a trained model in different contexts. To solve this limitation, we demonstrated the potential of using transfer learning. This technique enabled us to adapt the neural network representations learnt on a large source dataset to a different context by adjusting the pre-trained neural network weights using a limited amount data from the new context. Although the performance of the fine-tuned model increases with the amount of the new data made available, we showed that predictions can be drastically improved using as little as 36 new data points across different experimental conditions and setups. Our results suggest that a two-step approach, where a small-scale experiment is performed to fine-tune a pre-trained model in a new context, before it is then used for actual design.

The transfer learning protocol used here employs a lowered learning rate. While this approach is appealing due to its simplicity, it could be refined by using more advanced methods from few-shot learning to decrease the amount of training data needed from the new context ^72^. The transfer learning step itself could also suggest the most informative sequences to characterize, using for example different acquisition functions to reduce epistemic uncertainty ^73^. Furthermore, the deep learning models we used only considered the 5’UTR sequence alone, ignoring surrounding sequence information (e.g., the downstream protein coding region). While it is remarkable that only a handful of sequences are capable of adapting such models to new contexts, implicitly integrating information from the surrounding sequence in the deep learning models could increase model generalisation.

Data-centric approaches to biological design are already revolutionising how we design experiments ^74;75^, proteins ^76;77^ and genetic parts/circuits ^41;47^. However, being able to reuse and re-purpose machine learning models will be vital for broadening their reach beyond the constrained experiments used for training. The work presented here offers a practical approach to overcoming this issue and we hope will support the development of “context-aware” deep learning models tailored to the context in which they are being used to accelerate genetic circuit design workflows and reduce the need for trial-and-error when scaling the complexity of our biological designs.

## METHODS

### Dataset preparation

We inferred the log-normal distribution parameters for each sequence in the Kuo ^19^, Kosuri ^4^ and Evfratov ^64^ datasets using the FORECAST package ^51^ and raw sequencing data. The maximum fluorescence of the last sorting bin was set at 10^5^. The precision associated with each dataset was assessed using the width of the median 99.7% confidence interval obtained during the inference procedure. The median was used instead of the mean to avoid the outliers displaying abnormally large confidence intervals. Except for the Evfratov dataset, only constructs sorted into multiple bins and with an invertible hessian at the maximum likelihood estimates were kept. We did not perform inference on the Bonde dataset as we could not access the raw Flow-seq sequencing data. Instead, we relied on the values provided in the original paper ^68^. We also used the values from the original paper for the Hollerer dataset ^48^. For the Bonde, Kuo and Hollerer datasets, a train/validation/test split with ratios 80%/10%/10% was conducted using stratifying sampling with respect to the mean output fluorescence. For the Kosuri dataset, we filtered out the RBSs used to train the RBS Calculator ^21^ to avoid unfair comparisons. We discarded the promoters left with less than 60 RBSs after the whole filtering step. For each of the 63 promoters left, we assembled the corresponding test set by first selecting 30 sequences, with balanced log-fluorescence mean, and split the remaining sequences in the validation set (always 20 sequences) and the train set (ranging from 12 to 61 sequences according to the promoter). The extent of RBS strength spread for each dataset was robustly assessed by subtracting the 1st to the 99th quantile. This interval was then divided into 8 to define the margin of error. Following this, the accuracy was measured by the percentage of predictions within the margin of error centered around the ground truth. The width of this margin of error was 1.12 for the *fepB* dataset, 1.06 for the *arti* dataset, 1.28 for the *dmsC* dataset, 0.22 for the Bonde dataset and 1.6 for the Kosuri dataset.

### Activity cliff labeling

Test sequences from the *fepB* dataset were partitioned into two groups based on their log-fluorescence mean. Sequences below a value of 3 were classified as weak RBSs and sequences above a value of 6 were classified as strong RBSs. We then processed the weak and strong RBSs and labeled as activity cliffs the pairs of sequences differing by only mutation.

### Architecture considered, model training and hyper-parameter optimisation

Each model used as input the one-hot encoded 5’UTR sequence, followed by a particular backbone (LSTM, CNN, Conv+LSTM), resulting in an output matrix that was flattened and followed by a dense multi-linear perceptron (MLP) to predict the target output. For the Conv+LSTM model, the output of the Convolutional layer was a matrix of size [features, step] which was used as input for the LSTM layer. Models predicting the mean log-normal fluorescence distribution for each 5’UTR were trained by minimising the mean squared loss. We optimised the batch size, learning rate and architecture parameters for each candidate model (CNN or Conv-LSTM) to minimise the loss on the validation dataset. Learning rate was updated according to the ReduceLROnPlateau scheduler ^78^ with parameters mode = ‘min’, factor = 0.5, patience = 3, threshold = 0.01, min lr = 10^*−*5^. Models were trained for a maximum of 20 epochs and the loss on the validation dataset was monitored in order to use early stopping with a patience of 5 epochs to prevent over-fitting. Hyperparameters for the CNN and Conv-LSTM models trained on the *fepB* dataset were optimised using the Tree-structure Parzen estimator implemented in Optuna ^79^. We used a budget of 200 trials to optimise each architecture. Unpromising trials were pruned based on the intermediate value of the loss on the validation dataset. Parameter search space and optimal values for each architecture are noted in **Supplementary Table 1**.

### Scaling experiments

To assess the impact of the training dataset size on the model performance, we trained the Conv-LSTM model with the optimal hyper-parameters on a subset of the training and validation data. The test set sizedid not vary. The total number of training sequences took values in: 40, 50, 60, 70, 80, 90, 200, 500, 1,000, 2,000, 5,000, 10,000, 20,000, 50,000, 100,000, 120,000 with 90% of the examples for the training set and the remaining 10% for the validation set. For the smaller Bonde dataset, we limited the range of values to 2,000, which roughly correspond to the maximum training set size (2,487 examples in the training set). Experiments were conducted three times for each dataset with different seeds (3, 7, 13) to allow for various dataset splits and different initialisations of neural network parameters.

### Predicting standard deviation of log-normal distributions

The same Conv-LSTM backbone was used to model the distribution parameters for each sequence in the *fepB* dataset with the only difference being that the output layer included an additional neuron to capture the standard deviation log*s* of the log-normal fluorescence distribution. We used the Kullback-Leibler divergence ^80^ between the predicted and ground truth distribution as a loss function to train the model, following previous work ^58^.

### Transfer learning

Two transfer learning procedures were considered to fine-tune each pre-trained model for new contexts. The main one consisted of using each pre-trained model and further training it on the new dataset using the same mean squared loss, however, with a lower learning rate. All neural networks parameters could be updated. We used the ADAM optimizer with a learning rate of 3 *×* 10^*−*5^ and a batch size of 32 for a maximum of 600 epochs. The learning rate was updated according to the ReduceLROnPlateau scheduler ^78^ with parameters mode = ‘min’, factor = 0.5, patience = 3, threshold = 0.01, min lr = 10^*−*5^. We monitored the loss on the validation dataset and used early stopping with a patience of 80 epochs to prevent over-fitting. For the second transfer learning approach, we froze all parameters of the backbone Conv-LSTM model up until the last multi-layer perceptron. The perceptron for this model consisted of one intermediate layer with 128 nodes and one output node. Only parameters in the multi-layer perceptron were updated during the fine-tuning protocol, using the same approach as described above.

### Evaluation of fine-tuning for the Kosuri dataset

The Kosuri test set is small (30 samples for each promoter subset), which prevents accurate estimation of metrics. Therefore, we used the following pipeline to quantify and account for the measurement uncertainty. The standard error for the accuracy metric was estimated via 500 bootstrap samples of the test data using the quantile method. To detect statistical difference before and after fine-tuning, we conducted a paired permutation test on each promoter dataset. The null hypothesis was that fine-tuning does not have any effect on improving protein expression prediction from the sequence. The permutation test function from the MLXTEND package ^81^ was used with 10,000 permutations to allow for the detection of p-values as small as 10^*−*4^.

### RNA Foundational Model

We used the RNA foundational model (RNA-FM) developed by Chen and colleagues ^70^, which is based on the BERT bidirectional transformer architecture ^82^, trained on RNA-central ^83^ using a masked language modeling self-supervised learning scheme ^82^. We used this model to generate embeddings of shape (L, 640) for each 5’UTR sequence of length L. These embeddings were then used as inputs for a downstream architecture comprising 6 residual blocks of 1D convolutional layers followed by a flattening layer and a regression head to predict the mean log-fluorescence. This architecture was trained and fine-tuned on different datasets according to the same protocol described above.

### Alternative non-deep learning models

The RBS calculator v2.1 web server was used to compute the maximum translation initiation rate (TIR max) for each activity cliff in the *fepB* test set. However, using the web server to compute the TIR max is not amenable for larger test sets (typically *>*10,000 sequences). Therefore, we resorted to different non-deep learning based models to assess the deep learning model predictions for the *arti* and *dmsC* test datasets. Amongst them were the anti-Shine Dalgarno:Shine Dalgarno (aSD:SD) base pairing free energy and local mRNA folding free energy, which were extracted from the Kuo dataset ^19^ for each 5’UTR. EMOPEC and RBSeval models were also considered, with the predictions obtained using the implementation from Terai and colleagues ^69^.

### General computational tools

All analysis scripts and the deep learning pipeline were written in Python version 3.9 using numpy version 1.19.15^84^ for matrix algebra, scipy version 1.4.1 for statistical functions ^85^, Pytorch version 1.12^78^ for deep learning. Statistical inference speed comparisons between MOM and ML approaches were conducted on an Apple iMac computer with a 3 GHz 6-core Intel Core i5 processor and 16 GB of RAM.

## Data Availability

Code for the deep neural networks and transfer learning pipeline used in this study are available at: https://gitlab.com/Pierre-Aurelien/rebeca.

## ACKNOWLEDGMENTS

This work was supported by the EPSRC/BBSRC Centre for Doctoral Training in Synthetic Biology grant EP/L016494/1 (P.-A.G.), BrisEngBio, a UKRI-funded Engineering Biology Research Centre grant BB/W013959/1 (T.E.G.), a Turing Fellowship from The Alan Turing Institute under EPSRC grant EP/N510129/1 (T.E.G.), and a Royal Society University Research Fellowship grants UF160357 and URF*\*R*\*221008 (T.E.G.) The funders had no role in study design, data collection and analysis, decision to publish or preparation of the manuscript.

## AUTHOR CONTRIBUTIONS

P.-A.G. developed the methodology, carried out all computational experiments and performed the analysis. T.E.G. supervised the work and secured funding. Both authors conceived of the study and wrote the manuscript.

## TABLE OF CONTENTS FIGURE

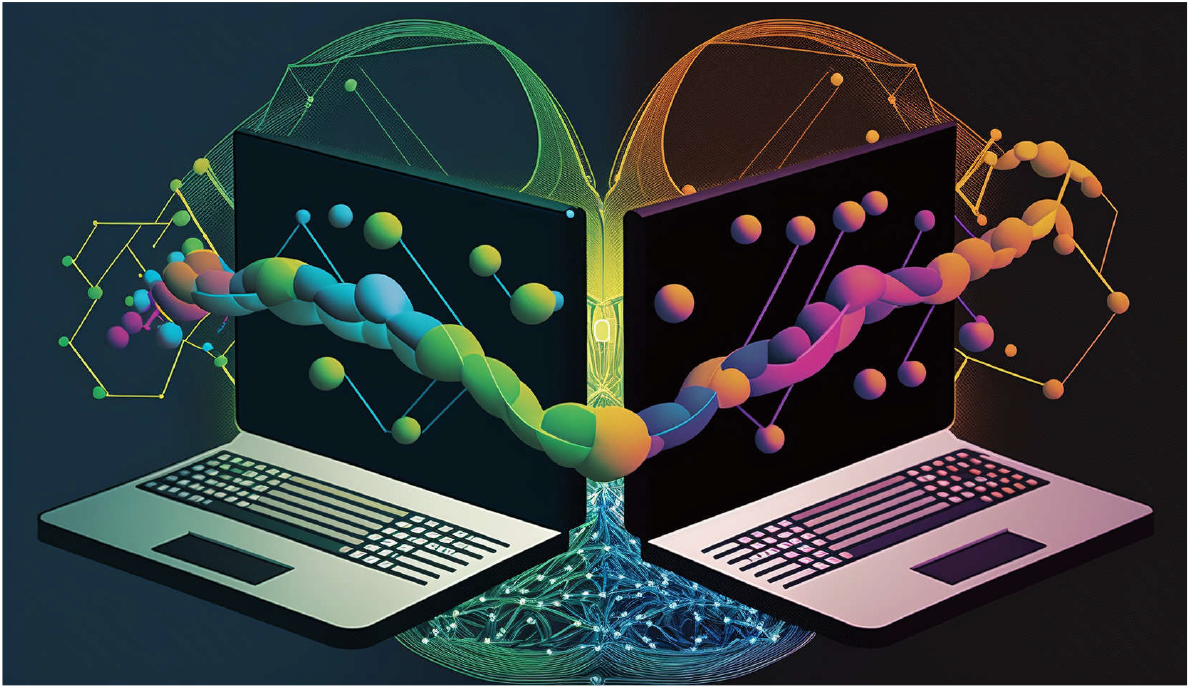

## Notes

### Competing Interest Statement

The authors have declared no competing interest.

## REFERENCES

[1] Nielsen, A. A. K. et al. Genetic circuit design automation. Science 352, aac7341 (2016).

[2] Cardinale, S. & Arkin, A. P. Contextualizing context for synthetic biology – identifying causes of failure of synthetic biological systems. Biotechnology Journal 7, 856–866 (2012).

[3] Yeung, E. et al. Biophysical Constraints Arising from Compositional Context in Synthetic Gene Networks. Cell Systems 5, 11–24.e12 (2017).

[4] Kosuri, S. et al. Composability of regulatory sequences controlling transcription and translation in Escherichia coli. Proceedings of the National Academy of Sciences 110, 14024–14029 (2013).

[5] Ceroni, F., Furini, S., Stefan, A., Hochkoeppler, A. & Giordano, E. A Synthetic Post-transcriptional Controller To Explore the Modular Design of Gene Circuits. ACS Synthetic Biology 1, 163–171 (2012).

[6] Frei, T. et al. Characterization and mitigation of gene expression burden in mammalian cells. Nature Communications 11, 4641 (2020).

[7] Jayanthi, S., Nilgiriwala, K. S. & Del Vecchio, D. Retroactivity Controls the Temporal Dynamics of Gene Transcription. ACS Synthetic Biology 2, 431–441 (2013).

[8] Kim, K. H. & Sauro, H. M. Fan-out in gene regulatory networks. Journal of Biological Engineering 4, 16 (2010).

[9] Borkowski, O., Ceroni, F., Stan, G.-B. & Ellis, T. Overloaded and stressed: Whole-cell considerations for bacterial synthetic biology. Current Opinion in Microbiology 33, 123–130 (2016).

[10] Cardinale, S., Joachimiak, M. P. & Arkin, A. P. Effects of Genetic Variation on the E. coli Host-Circuit Interface. Cell Reports 4, 231–237 (2013).

[11] Dahl, R. H. et al. Engineering dynamic pathway regulation using stress-response promoters. Nature Biotechnology 31, 1039–1046 (2013).

[12] Chory, E. J., Gretton, D. W., DeBenedictis, E. A. & Esvelt, K. M. Enabling high-throughput biology with flexible open-source automation. Molecular Systems Biology 17, e9942 (2021).

[13] Johns, N. I. et al. Metagenomic mining of regulatory elements enables programmable species-selective gene expression. Nature Methods 15, 323–329 (2018).

[14] Moser, F. et al. Genetic Circuit Performance under Conditions Relevant for Industrial Bioreactors. ACS Synthetic Biology 1, 555–564 (2012).

[15] Gorochowski, T. E., van den Berg, E., Kerkman, R., Roubos, J. A. & Bovenberg, R. A. L. Using synthetic biological parts and microbioreactors to explore the protein expression characteristics of escherichia coli. ACS Synthetic Biology 3, 129–139 (2014). PMID: 24299494.

[16] Beal, J. et al. Quantification of bacterial fluorescence using independent calibrants. PLOS ONE 13, 1–15 (2018).

[17] Marguet, P., Balagadde, F., Tan, C. & You, L. Biology by design: Reduction and synthesis of cellular components and behaviour. Journal of The Royal Society Interface 4, 607–623 (2007).

[18] Tietze, L. & Lale, R. Importance of the 5′ regulatory region to bacterial synthetic biology applications. Microbial Biotechnology 14, 2291–2315 (2021).

[19] Kuo, S.-T. et al. Global fitness landscapes of the Shine-Dalgarno sequence. Genome Research (2020).

[20] Egbert, R. G. & Klavins, E. Fine-tuning gene networks using simple sequence repeats. Proceedings of the National Academy of Sciences 109, 16817–16822 (2012).

[21] Salis, H. M., Mirsky, E. A. & Voigt, C. A. Automated design of synthetic ribosome binding sites to control protein expression. Nature Biotechnology 27, 946–950 (2009).

[22] Goodman, D. B., Church, G. M. & Kosuri, S. Causes and Effects of N-Terminal Codon Bias in Bacterial Genes. Science (2013).

[23] Kudla, G., Murray, A. W., Tollervey, D. & Plotkin, J. B. Coding-Sequence Determinants of Gene Expression in Escherichia coli. Science 324, 255–258 (2009).

[24] Gorochowski, T. E., Ignatova, Z., Bovenberg, R. A. & Roubos, J. A. Trade-offs between tRNA abundance and mRNA secondary structure support smoothing of translation elongation rate. Nucleic Acids Research 43, 3022–3032 (2015).

[25] Wohlgemuth, S. E., Gorochowski, T. E. & Roubos, J. A. Translational sensitivity of the Escherichia coli genome to fluctuating tRNA availability. Nucleic Acids Research 41, 8021–8033 (2013).

[26] Del Vecchio, D. Modularity, context-dependence, and insulation in engineered biological circuits. Trends in Biotechnology 33, 111–119 (2015).

[27] Mutalik, V. K. et al. Precise and reliable gene expression via standard transcription and translation initiation elements. Nature Methods 10, 354–360 (2013).

[28] Carr, S. B., Beal, J. & Densmore, D. M. Reducing dna context dependence in bacterial promoters. PLOS ONE 12, 1–15 (2017).

[29] Park, Y., Espah Borujeni, A., Gorochowski, T. E., Shin, J. & Voigt, C. A. Precision design of stable genetic circuits carried in highly-insulated e. coli genomic landing pads. Molecular Systems Biology 16, e9584 (2020).

[30] Qi, L., Haurwitz, R. E., Shao, W., Doudna, J. A. & Arkin, A. P. Rna processing enables predictable programming of gene expression. Nature Biotechnology 30, 1002–1006 (2012).

[31] Lou, C., Stanton, B., Chen, Y.-J., Munsky, B. & Voigt, C. A. Ribozyme-based insulator parts buffer synthetic circuits from genetic context. Nature Biotechnology 30, 1137–1142 (2012).

[32] Davis, J. H., Rubin, A. J. & Sauer, R. T. Design, construction and characterization of a set of insulated bacterial promoters. Nucleic Acids Research 39, 1131–1141 (2011).

[33] Mishra, D., Rivera, P. M., Lin, A., Del Vecchio, D. & Weiss, R. A load driver device for engineering modularity in biological networks. Nature Biotechnology 32, 1268–1275 (2014).

[34] Ceroni, F. et al. Burden-driven feedback control of gene expression. Nature Methods 15, 387–393 (2018).

[35] Jones, R. D. et al. An endoribonuclease-based feedforward controller for decoupling resource-limited genetic modules in mammalian cells. Nature Communications 11, 5690 (2020).

[36] Del Vecchio, D., Ninfa, A. J. & Sontag, E. D. Modular cell biology: Retroactivity and insulation. Molecular Systems Biology 4, 161 (2008).

[37] Bartoli, V., Meaker, G. A., di Bernardo, M. & Gorochowski, T. E. Tunable genetic devices through simultaneous control of transcription and translation. Nature Communications 11, 2095 (2020).

[38] Bartoli, V., di Bernardo, M. & Gorochowski, T. E. Self-adaptive biosystems through tunable genetic parts and circuits. Current Opinion in Systems Biology 24, 78–85 (2020). Systems immunology hostpathogen interaction (2020).

[39] Tas, H., Grozinger, L., Stoof, R., de Lorenzo, V. & Goni-Moreno, A. Contextual dependencies expand the re-usability of genetic inverters. Nature Communications 12, 355 (2021).

[40] Castle, S. D., Grierson, C. S. & Gorochowski, T. E. Towards an engineering theory of evolution. Nature Communications 12, 3326 (2021).

[41] Cambray, G., Guimaraes, J. C. & Arkin, A. P. Evaluation of 244,000 synthetic sequences reveals design principles to optimize translation in Escherichia coli. Nature Biotechnology 36, 1005–1015 (2018).

[42] Tarnowski, M. J. & Gorochowski, T. E. Massively parallel characterization of engineered transcript isoforms using direct RNA sequencing. Nature Communications 13, 434 (2022).

[43] Cadwell, R. C. & Joyce, G. F. Randomization of genes by PCR mutagenesis. Genome Research 2, 28–33 (1992).

[44] Vidal, L. S., Isalan, M., Heap, J. T. & Ledesma-Amaro, R. A primer to directed evolution: Current methodologies and future directions. RSC Chemical Biology (2023).

[45] Ellis, T., Wang, X. & Collins, J. J. Diversity-based, model-guided construction of synthetic gene networks with predicted functions. Nature Biotechnology 27, 465–471 (2009).

[46] de Boer, C. G. et al. Deciphering eukaryotic gene-regulatory logic with 100 million random promoters. Nature Biotechnology 38, 56–65 (2020).

[47] Angenent-Mari, N. M., Garruss, A. S., Soenksen, L. R., Church, G. & Collins, J. J. A deep learning approach to programmable RNA switches. Nature Communications 11, 5057 (2020).

[48] Höllerer, S. et al. Large-scale DNA-based phenotypic recording and deep learning enable highly accurate sequence-function mapping. Nature Communications 11, 3551 (2020).

[49] Valeri, J. A. et al. Sequence-to-function deep learning frameworks for engineered riboregulators. Nature Communications 11, 5058 (2020).

[50] Sample, P. J. et al. Human 5′ UTR design and variant effect prediction from a massively parallel translation assay. Nature Biotechnology 37, 803–809 (2019).

[51] Gilliot, P.-A. & Gorochowski, T. E. Effective design and inference for cell sorting and sequencing based massively parallel reporter assays (2022).

[52] Nikolados, E.-M., Wongprommoon, A., Aodha, O. M., Cambray, G. & Oyarzún, D. A. Accuracy and data efficiency in deep learning models of protein expression. Nature Communications 13, 7755 (2022).

[53] Yosinski, J., Clune, J., Bengio, Y. & Lipson, H. How transferable are features in deep neural networks? In Advances in Neural Information Processing Systems, vol. 27 (Curran Associates, Inc., 2014).

[54] Liu, Z. et al. A ConvNet for the 2020s (2022). arXiv:2201.03545.

[55] Deng, J. et al. ImageNet: A large-scale hierarchical image database. In 2009 IEEE Conference on Computer Vision and Pattern Recognition, 248–255 (2009).

[56] Eraslan, G., Avsec, Ž., Gagneur, J. & Theis, F. J. Deep learning: New computational modelling techniques for genomics. Nature Reviews Genetics 20, 389–403 (2019).

[57] Beal, J. Biochemical complexity drives log-normal variation in genetic expression. Engineering Biology 1, 55–60 (2017).

[58] Kingma, D. P. & Welling, M. Auto-Encoding Variational Bayes. arXiv:1312.6114 [cs, stat] (2013). 1312.6114.

[59] Pfleger, B. F., Pitera, D. J., Smolke, C. D. & Keasling, J. D. Combinatorial engineering of intergenic regions in operons tunes expression of multiple genes. Nature Biotechnology 24, 1027–1032 (2006).

[60] Wang, H. H. et al. Programming cells by multiplex genome engineering and accelerated evolution. Nature 460, 894–898 (2009).

[61] Meng, H. et al. Quantitative Design of Regulatory Elements Based on High-Precision Strength Prediction Using Artificial Neural Network. PLOS ONE 8, e60288 (2013).

[62] Espah Borujeni, A., Channarasappa, A. S. & Salis, H. M. Translation rate is controlled by coupled trade-offs between site accessibility, selective RNA unfolding and sliding at upstream standby sites. Nucleic Acids Research 42, 2646–2659 (2014).

[63] Osterman, I. A., Evfratov, S. A., Sergiev, P. V. & Dontsova, O. A. Comparison of mRNA features affecting translation initiation and reinitiation. Nucleic Acids Research 41, 474–486 (2013).

[64] Evfratov, S. A. et al. Application of sorting and next generation sequencing to study 5’-UTR influence on translation efficiency in Escherichia coli. Nucleic Acids Research 45, 3487–3502 (2017).

[65] Breiman, L. Bagging predictors. Machine Learning 24, 123–140 (1996).

[66] Fan, H. et al. Transcription–translation coupling: Direct interactions of RNA polymerase with ribosomes and ribosomal subunits. Nucleic Acids Research 45, 11043–11055 (2017).

[67] Bakshi, S., Choi, H. & Weisshaar, J. C. The spatial biology of transcription and translation in rapidly growing Escherichia coli. Frontiers in Microbiology 6 (2015).

[68] Bonde, M. T. et al. Predictable tuning of protein expression in bacteria. Nature Methods 13, 233–236 (2016).

[69] Terai, G. & Asai, K. Improving the prediction accuracy of protein abundance in Escherichia coli using mRNA accessibility. Nucleic Acids Research 48, e81 (2020).

[70] Chen, J. et al. Interpretable RNA Foundation Model from Unannotated Data for Highly Accurate RNA Structure and Function Predictions (2022). arXiv:2204.00300.

[71] Vaswani, A. et al. Attention Is All You Need (2017). arXiv:1706.03762.

[72] Finn, C., Abbeel, P. & Levine, S. Model-Agnostic Meta-Learning for Fast Adaptation of Deep Networks. In Proceedings of the 34th International Conference on Machine Learning, 1126–1135 (PMLR, 2017).

[73] Gal, Y., Islam, R. & Ghahramani, Z. Deep Bayesian Active Learning with Image Data (2017). arXiv: 1703.02910.

[74] Pandi, A. et al. A versatile active learning workflow for optimization of genetic and metabolic networks. Nature Communications 13, 3876 (2022).

[75] Radivojević, T., Costello, Z., Workman, K. & Garcia Martin, H. A machine learning Automated Recommendation Tool for synthetic biology. Nature Communications 11, 4879 (2020).

[76] Anishchenko, I. et al. De novo protein design by deep network hallucination. Nature 600, 547–552 (2021).

[77] Tack, D. S. et al. The genotype-phenotype landscape of an allosteric protein. Molecular Systems Biology 17, e10179 (2021).

[78] Paszke, A. et al. PyTorch: An Imperative Style, High-Performance Deep Learning Library (2019). arXiv:1912.01703.

[79] Akiba, T., Sano, S., Yanase, T., Ohta, T. & Koyama, M. Optuna: A Next-generation Hyperparameter Optimization Framework. In Proceedings of the 25th ACM SIGKDD International Conference on Knowledge Discovery & Data Mining, KDD ‘19, 2623–2631 (Association for Computing Machinery, New York, NY, USA, 2019).

[80] Robert, C. P. Intrinsic losses. Theory and Decision 40, 191–214 (1996).

[81] Raschka, S. MLxtend: Providing machine learning and data science utilities and extensions to Python’s scientific computing stack. Journal of Open Source Software 3, 638 (2018).

[82] Devlin, J., Chang, M.-W., Lee, K. & Toutanova, K. BERT: Pre-training of Deep Bidirectional Transformers for Language Understanding (2019). arXiv:1810.04805.

[83] RNAcentral Consortium. RNAcentral 2021: Secondary structure integration, improved sequence search and new member databases. Nucleic Acids Research 49, D212–D220 (2021).

[84] Harris, C. R. et al. Array programming with NumPy. Nature 585, 357–362 (2020).

[85] Virtanen, P. et al. SciPy 1.0: Fundamental algorithms for scientific computing in Python. Nature Methods 17, 261–272 (2020).

