## Supplementary Information for "Transfer learning for cross-context prediction of protein expression from 5’UTR sequence"

**Table 1.** Hyperparameter optimisation results for the *fepB* context. Range of all hyperparameters considered when optimising the Conv-LSTM and CNN models.

| Hyperparameter | Search Space | |
|---|---|---|
| | Conv-LSTM | CNN |
| CNN Channels | [64, 128, **256**, 512] | [64, 128, **256**, 512] |
| CNN Layers | [1, **2**, 3] | [1, 2, **3**] |
| CNN Kernel | [4, 6, **8**, 10] | [4, 6, 8, **10**] |
| CNN Pool | [**1**, 2] | [**1**, 2] |
| LSTM Hidden Size | [5, 10, 25, **50**, 100, 200, 500] | ∅ |
| BILSTM | [**True**, False] | ∅ |
| Activation | [ReLU, ELU, **LeakyReLU**] | [ReLU, ELU, **LeakyReLU**] |
| MLP Hidden Size | [64, **128**, 256, 512, 1024] | [64, 128, 256, 512, **1024**] |
| MLP Layers | [**1**, 2, 3] | [1, **2** ,3] |
| Dropout Rate | [0..**0.28**..0.5] | [0..**0.36**..0.5] |
| Learning Rate | [1e-4..**3e-4**..1e2] | [1e-4..**2.6e-4**..1e2] |
| Batch Size | [**32**, 64] | [32, **64**] |
| Number of parameters | 1, 180, 837 | 8, 659, 713 |

**Table 2.** Performance metrics for the *arti* context test set for several RBS strength prediction algorithms.

| Algorithm \ Metric | Pearson $R^2$ | Spearman $\rho$ | Kendall $\tau$ |
|---|---|---|---|
| EMOPEC | 0.50 | 0.35 | 0.24 |
| RBSeval | 0.58 | 0.62 | 0.44 |
| Neural Network (fepb trained, before fine-tuning) | 0.73 | 0.74 | 0.55 |
| $\Delta G$ SD:aSD base pairing | $-0.5$ | $-0.32$ | $-0.23$ |
| $\Delta G$ mRNA folding | $4 \times 10^{-3}$ | 0.12 | 0.08 |

**Table 3.** Performance metrics for the *dmsC* context test set for several RBS strength prediction algorithms.

| Algorithm \ Metric | Pearson $R^2$ | Spearman $\rho$ | Kendall $\tau$ |
|---|---|---|---|
| EMOPEC | 0.47 | 0.42 | 0.29 |
| RBSeval | 0.47 | 0.56 | 0.39 |
| Neural Network (fepb trained, before fine-tuning) | 0.60 | 0.62 | 0.45 |
| $\Delta G$ SD:aSD base pairing | −0.42 | −0.26 | −0.19 |
| $\Delta G$ mRNA folding | 0.01 | 0.15 | −0.11 |

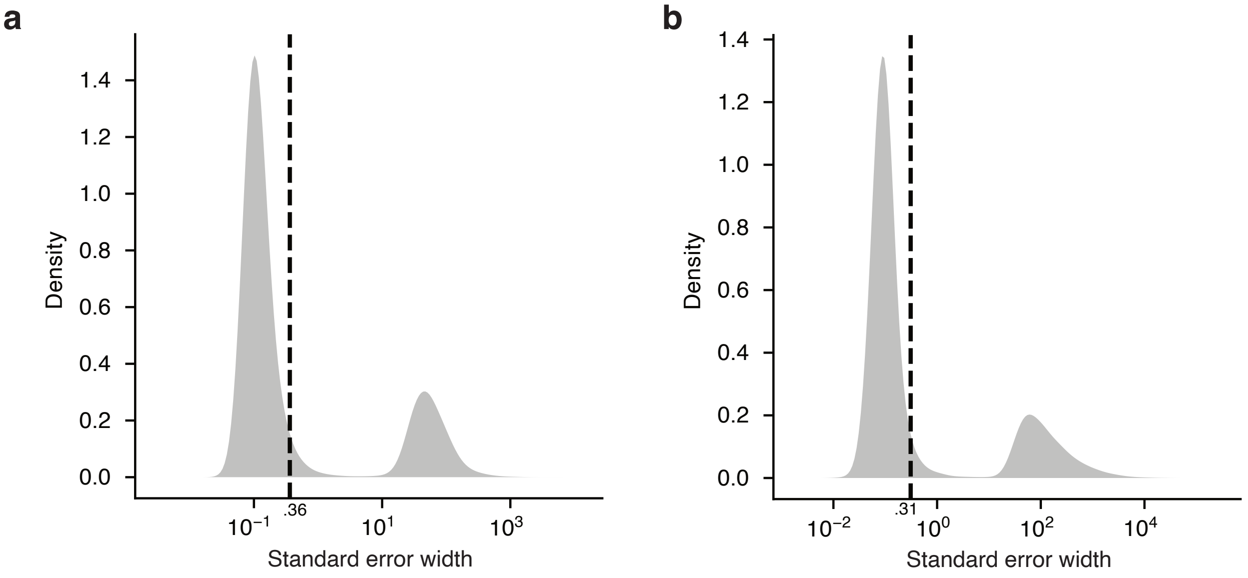

**Supplementary Figure 1: Precision of Flow-seq estimates.** Distributions of the standard error width associated with the Maximum likelihood estimates of the (**a**) fluorescence mean and (**b**) standard deviation for each sequence from the *fepB* context of the Kuo experiment. Dashed line indicates the median 99.7 % confidence interval for each statistic (0.36 for the log-fluorescence mean and 0.31 for the log-fluorescence standard deviation.)

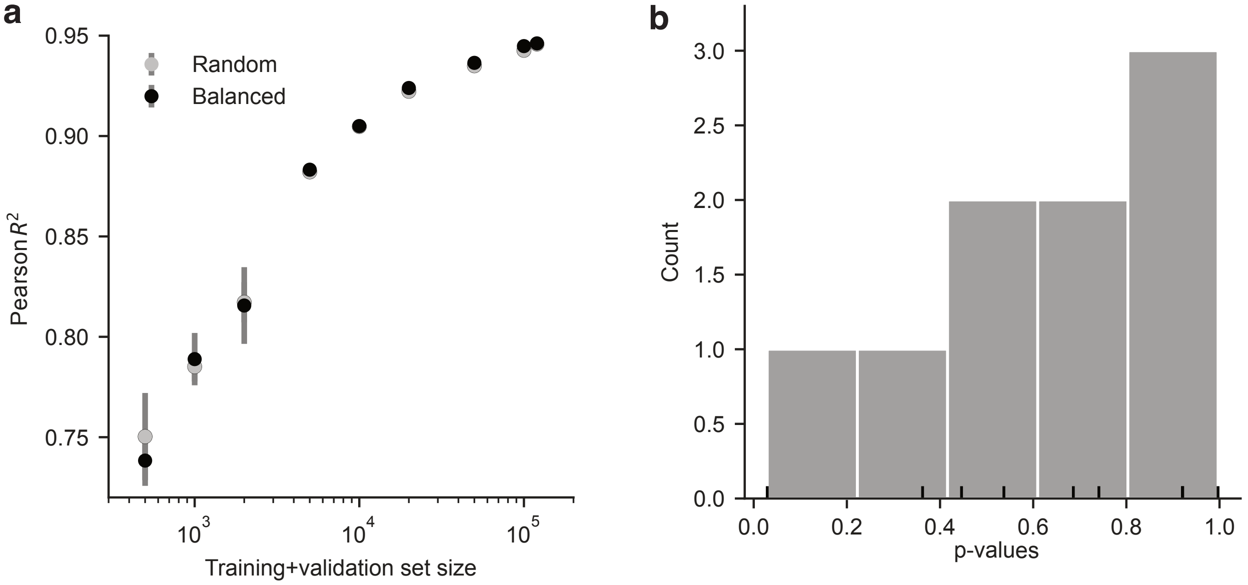

**Supplementary Figure 2: Accounting for the uncertainty of the neural network random initialisation.** (**a**) Performance of the best Conv-LSTM model predicting the log-fluorescence mean using varying size and composition of training data (*fepB* context). Pearson $R^2$ coefficient was evaluated on the test set when using a training set of sequences randomly selected (random) or selected to obtain a balanced spread of log-fluorescence mean values (balanced). Error bars indicate the standard deviation observed when training three models using different random seeds. (**b**) Distribution of the *p*-values obtained using multiple Welsch t-tests to challenge the null hypothesis of no effect observed when training a neural network on random vs balanced dataset.

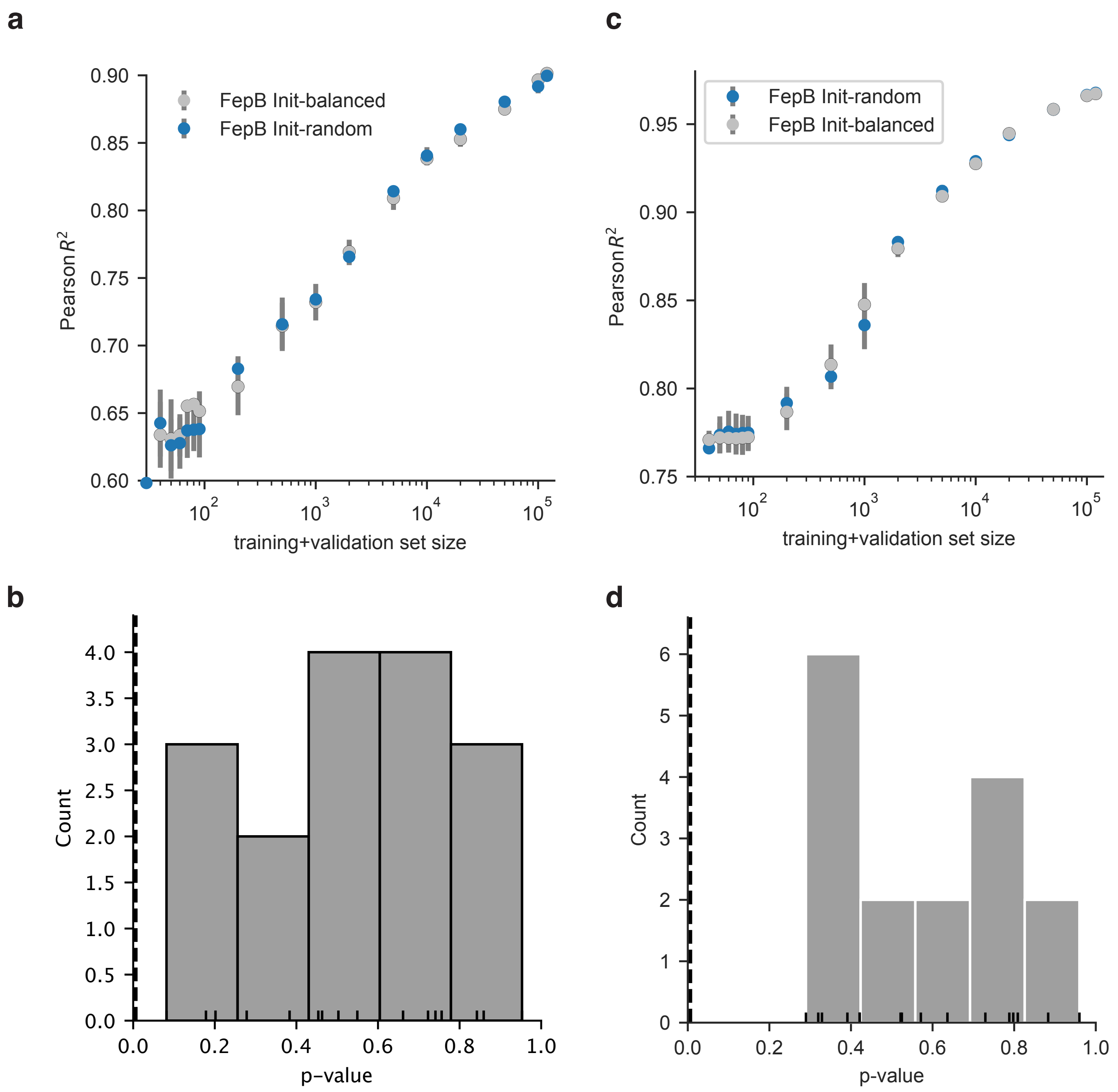

**Supplementary Figure 3: Stratified sampling of sequences do not improve fine-tuning.** (**a**) Test set performance (*dmsc* context of the Kuo dataset) of the Conv-LSTM model pre-trained on *fepB* context and fine-tuned on *dmsC*. (**b**) Distribution of p-values testing the statistical difference between the mean of the Pearson $R^2$, evaluated on the *dmsC* test set, between the random vs stratified sampling treatment. p-values are indicated with a tick on the $x$-axis. Threshold in dash is equal to $0.05/16 = 3 \times 10^{-3}$ (**c**) Test set performance (*arti* context of the Kuo dataset) of the Conv-LSTM model pre-trained on *fepB* context and fine-tuned on *arti*. (**d**) Distribution of p-values testing the statistical difference between the mean of the Pearson $R^2$, evaluated on the *arti* test set, between the random vs stratified sampling treatment. p-values are indicated with a tick on the $x$-axis. Threshold in dash is equal to was set to $0.05/16 = 3 \times 10^{-3}$

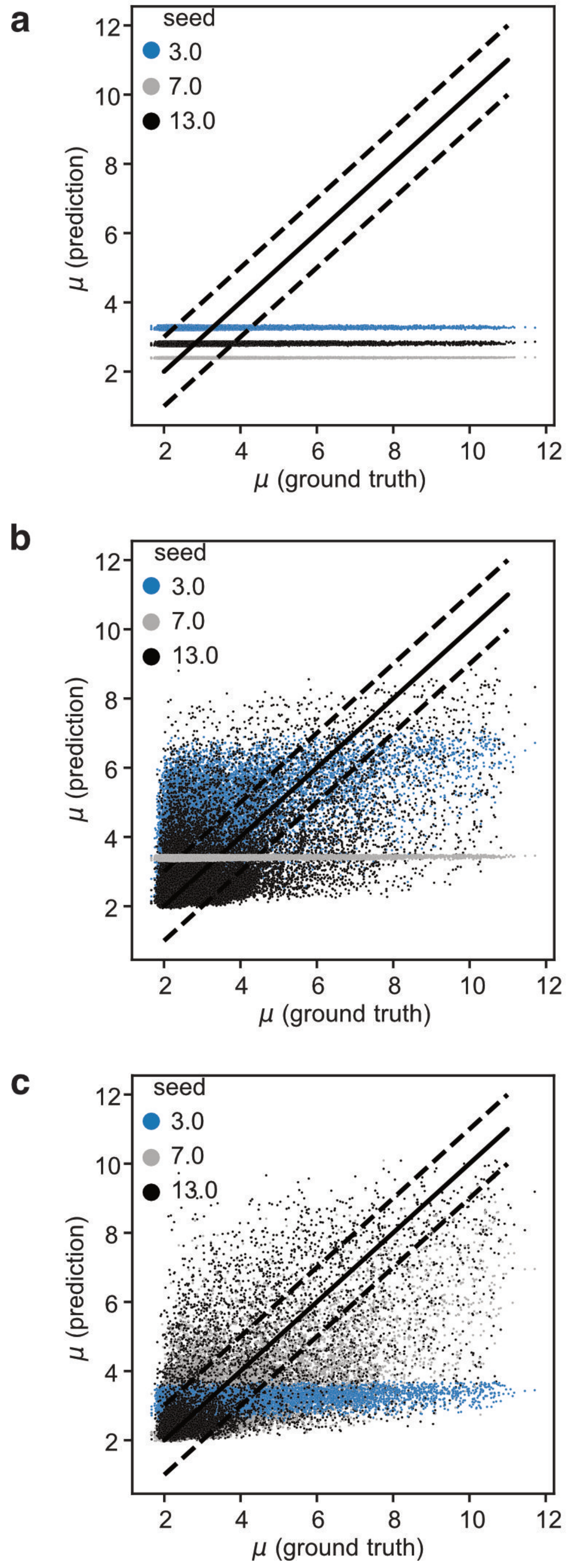

**Supplementary Figure 4: Visualising *arti* test set predictions.** Prediction of the hybrid Conv-LSTM neural network trained from scratch on the *arti* dataset with varying amount of examples in the training and validation set: (**a**) 90, (**b**) 500 and (**c**) 1000. Conv-LSTM training was carried out in triplicates using different random seeds (3, 7 and 13, respectively).

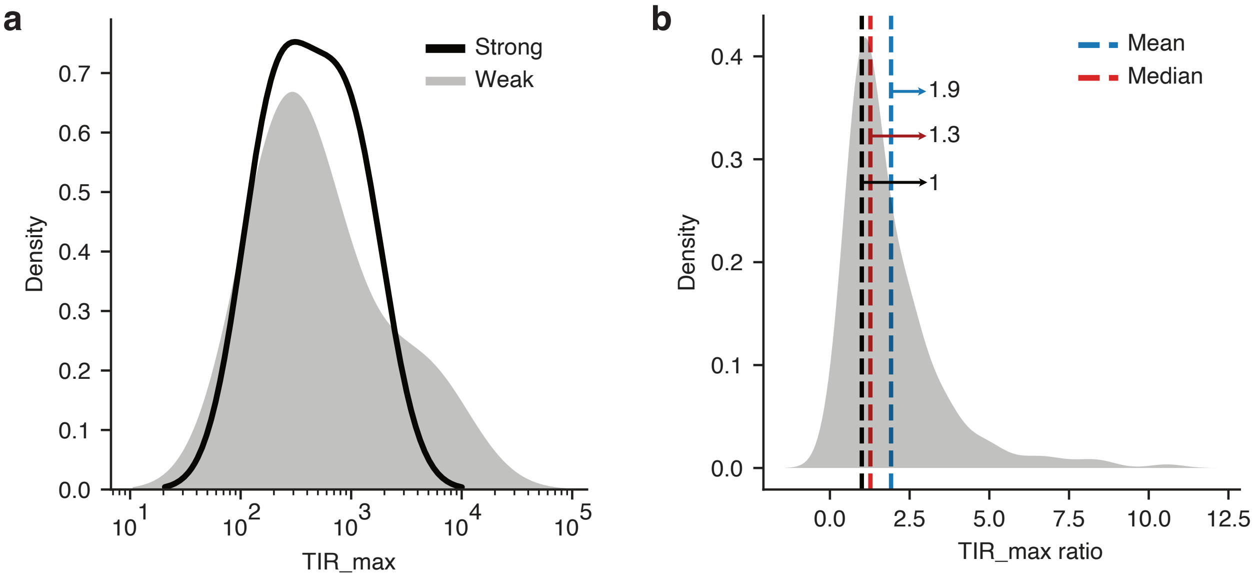

**Supplementary Figure 5: Predictions on activity cliffs using Salis RBS calculator v2.1 .** a) Distributions of maximum translation initiation rates (TIR_max) using the RBS calculator for the weak (mean log fluorescence $\mu < 3$) and strong (mean log-fluorescence $\mu > 6$) 5'UTR sequences from the *fepB* test set. b) Distribution of the TIR_max ratio between pairs of activity cliffs from the *fepB* test set. The ratio was taken between the TIR_max of the strong 5'UTR and the TIR_max of the weak 5'UTR sequence.Both sequences only differ by only one mutation.

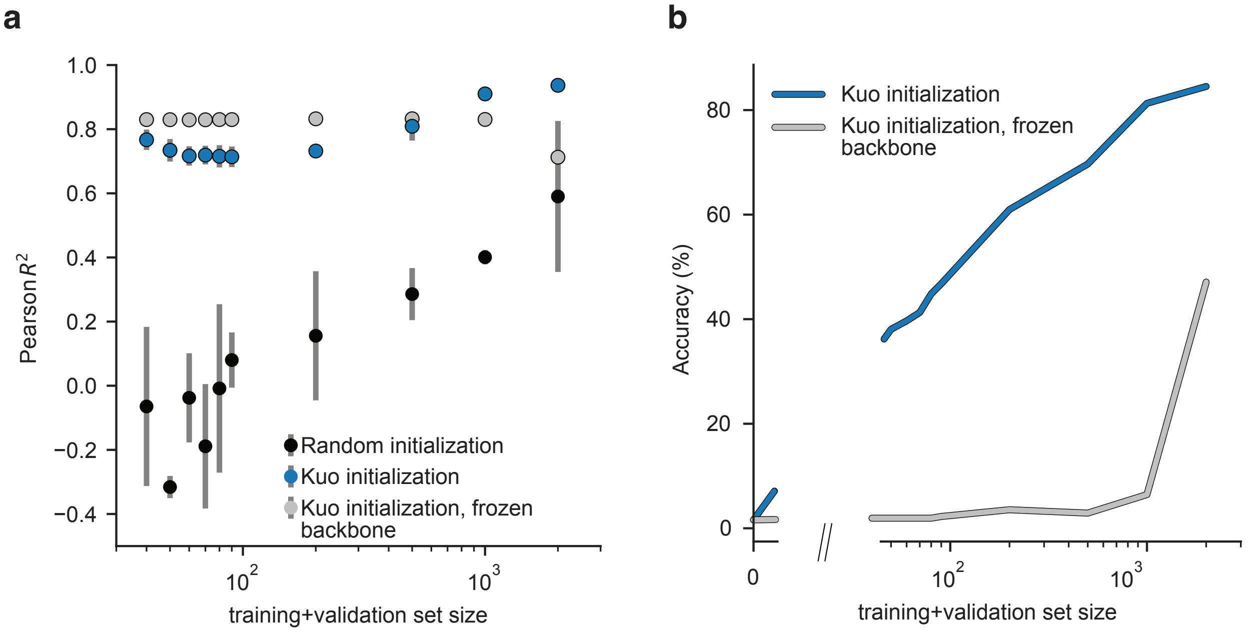

**Supplementary Figure 6: Comparing fine-tuning procedures on Bonde dataset.** (**a**) Performance on the Bonde dataset of the hybrid Conv-LSTM model trained from scratch (Random initialization) or by fine-tuning a model pre-trained on the Kuo dataset. Fine-tuning was accomplished by updating all model parameters (Kuo initialization) or with frozen backbone parameters (Kuo initialization, frozen backbone). (**a**) Performance on the Bonde dataset of the hybrid Conv-LSTM pre-trained on the Kuo dataset and fine-tuned by either updating all model parameters (Kuo initialization) or with frozen backbone parameters (Kuo initialization, frozen backbone).

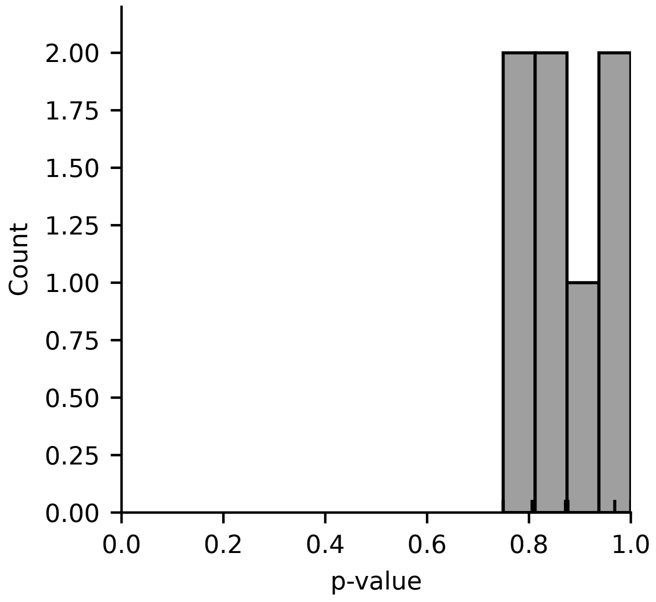

**Supplementary Figure 7: Negative transfer on Kosuri dataset.** Distribution of p-values using paired permutation tests for equal mean before and after fine-tuning for promoter 0 to promoter 7 from the Kosuri dataset. The predictions come from the Conv-LSTM model trained on all the Kuo data.

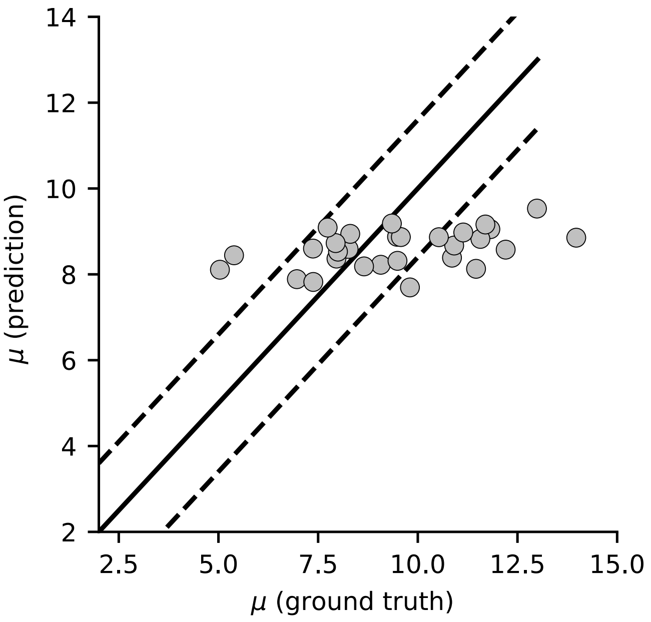

**Supplementary Figure 8: RBSeval gives uninformative predictions.** Distribution of p-values using paired permutation tests for equal mean after and before fine-tuning for promoter 0 to promoter 7 from the Kosuri dataset. The predictions come from the Conv-LSTM model trained on all the Kuo data.
